# Sindbis virus is suppressed in the yellow fever mosquito *Aedes aegypti* by ATG-6/Beclin-1 mediated activation of autophagy

**DOI:** 10.1101/2023.02.02.526867

**Authors:** Sujit Pujhari, Chan C. Heu, Marco Brustolin, Rebecca M. Johnson, Donghun Kim, Jason L. Rasgon

**Affiliations:** Department of Entomology, the Center for Infectious Disease Dynamics, and the Huck Institutes of the Life Sciences, Pennsylvania State University, University Park, PA, United States; Department of Pharmacology, Physiology and Neuroscience, School of Medicine, University of South Carolina, Columbia, SC; USDA-ARS, Maricopa, AZ, USA; Unit of Entomology, Department of Biomedical Sciences, Institute of Tropical Medicine, Antwerp, Belgium; Department of Entomology, The Connecticut Agricultural Experiment Station, New Haven, CT; Department of Vector Entomology, Kyungpook National University, Daegu, South Korea

## Abstract

Autophagy is a critical modulator of pathogen invasion response in vertebrates and invertebrates. However, how it affects mosquito-borne viral pathogens that significantly burden public health remains underexplored. To address this gap, we use a genetic approach to activate macroautophagy/autophagy in the yellow fever mosquito (*Aedes aegypti*), infected with a recombinant Sindbis virus (SINV) expressing an autophagy activator. We first demonstrate a 17- amino acid peptide derived from the *Ae. aegypti* autophagy-related protein 6 (ATG-6/beclin-1-like protein) is sufficient to induce autophagy in C6/36 mosquito cells, as marked by lipidation of ATG- 8 and puncta formation. Next, we engineered a recombinant SINV expressing this bioactive beclin- 1-like peptide and used it to infect and induce autophagy in adult mosquitoes. We find that modulation of autophagy using this recombinant SINV negatively regulated production of infectious viruses. The results from this study improve our understanding of the role of autophagy in arboviruses in invertebrate hosts and also highlight the potential for the autophagy pathway to be exploited for arboviral control.

## Introduction

Autophagy is a highly conserved catabolic pathway in eukaryotes that contributes to maintaining cellular homeostasis by providing a mechanism for protein and organelle quality control. ^1^ Autophagy plays a role in cellular defense against acute stress (and stress induced by nutritional starvation in particular), and is also involved in critical physiological and developmental processes. ^2,3-13^ Autophagy is a localized phenomenon where cytoplasmic components targeted for degradation are first sequestered into a double-membrane vesicle, digested by lysosomal enzymes to their functional units, then released back to the cytoplasm where they are available for maintaining basal cellular activities. ^1, 14 4-13^

The molecular regulators of autophagy have been well-characterized. Under normal, nutrition-rich condition, autophagy is inhibited by the TOR and other regulatory kinases. ^15^ However, in nutrient-poor conditions, the energy sensor AMPK (adenosine monophosphate-activated protein kinase) is activated, leading to the phosphorylation of the ATG1/ULK1 complex, which is the key initiator of autophagy. ^16^ Additionally, activated AMPK phosphorylates TSC2, which leads to the inhibition of TOR, a protein that acts as a negative regulator of autophagy, resulting in the activation of autophagic processes. In addition, this leads to the activation of the FOXO (forkhead box O) transcription factor that activates autophagy-related (ATG) genes and promotes the process of autophagy. ^17-21^ One key regulator of autophagy is the protein ATG-6, also known as beclin-1, which plays a crucial role in activating autophagy. ^22, 23^ ATG-8 (also known as LC3) is another important protein in the autophagy process and acts as a marker of autophagy, as it is conjugated to phosphatidylethanolamine and recruited to the phagophore membrane during autophagosome formation. ^24, 25^

Autophagy-related genes work in concert to orchestrate five distinct steps of autophagy: (1) initiation, (2) *de novo* formation of the phagophore at the endoplasmic reticulum or other membranes, (3) maturation or closure of the phagophore to generate the autophagosome, (4) fusion of the autophagosome with lysosomes and (5) cargo degradation and cytosolic recycling of metabolites. ^1^ Beclin-1 is a critical and rate-limiting protein for the initiation step of autophagy. It acts via its interaction with the class III phosphatidylinositol-3-kinase/Vps34 complex (VPS15- VPS34-ATG14-beclin1). ^26^ Inactivation or loss of the beclin-1 gene causes severe phenotypes across diverse animals, including cell apoptosis in *Caenorhabditis elegans* and increased aggregation and neurodegeneration in mice and humans. ^27, 28, 22 29^ In mosquitoes, autophagy plays a central role in the progression of gonadotropic cycles, where autophagy-incompetent females exhibit slow and abnormal egg maturation. ^29^

Autophagy is also involved in innate immunity against intracellular pathogens such as viruses ^4-13^, including clearance of SINV in mice. ^30-32^. However, some viruses subvert autophagy-based defenses and even hijack autophagic components to escape antiviral defense systems and increase their replication, assembly, and release. ^10, 19, 33-35^ For example, non-structural proteins NS4A and NS4B of Zika virus (ZIKV) and NS4A of Dengue virus (DENV) interact with Akt-mTOR in vertebrates to induce autophagy and support viral replication. ^30, 33-35^ In contrast, West Nile virus (WNV) capsid protein inhibits autophagy by downregulating AMP-activated protein kinase, which also results in neuropathogencity. ^19^ While this and other evidence highlight the importance of autophagy in host-virus dynamics, most of our understanding comes from studies in vertebrates, and interactions between arboviruses and autophagic pathways in invertebrate hosts remain poorly understood.

Here we address this knowledge gap by studying the role of autophagy in mosquito responses to infection with Sindbis virus (SINV); the prototype of the genus *Alphavirus* (family Togaviridae). SINV is a positive-sense single-stranded RNA virus with a broad host range that can be transmitted by the yellow fever mosquito *Aedes aegypti*. ^36-38^ We used this virus-mosquito model system to functionally test the effects of autophagy on arboviral infection. We engineered a recombinant SINV to express a 17-amino acid beclin-1-like peptide from *Ae. aegypti* that induces autophagy, then tracked SINV replication *in vitro* and *in vivo*. ^39^ Our results demonstrate that autophagy negatively impacts SINV replication *in vitro* and depresses SINV titers both *in vitro* and *in vivo* in *Ae. aegypti* mosquitoes.

## Results

### *Ae. aegypti* beclin-1-like peptide is conserved

To study beclin-1-like protein in *Ae. aegypti*, we analyzed and compared this protein sequence to its human orthologue and invertebrate homologs (*Anopheles gambiae, Drosophila melanogaster*, and *Caenorhabditis elegans)*. Phylogenetically, the insect sequences clustered together while *C. elegens* beclin-1 sequence was most diverse (Figures 1A and S1). *Ae. aegypti* beclin-1 is 426 aa long, compared to the 450 aa human orthologue. The C-terminus of these two protein sequences is highly conserved with 77% sequence identity. We noted, in particular, a 17-amino acid peptide of beclin-1 (243-260 aa in *Ae. aegypti*) that is highly conserved across these taxa. Using human beclin-1 as the template (PDB id 4ddp) we generated a 3D model of the *Ae. aegypti* beclin-1. This showed that the conserved 17-aa peptide is located on the surface of the protein (Figure 1B).

**Figure 1.**
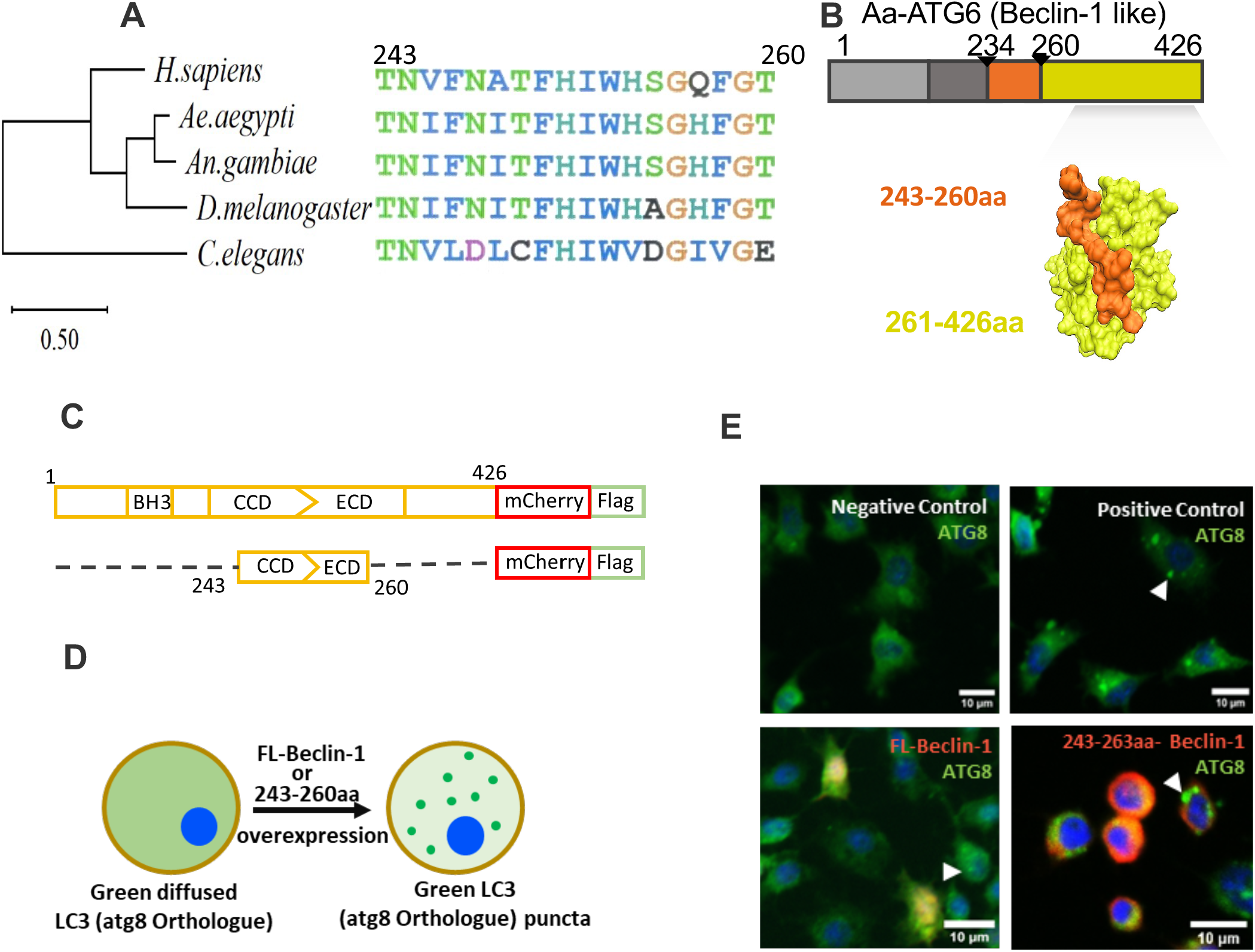
A 17-amino acid peptide of *Aedes aegypti* ATG6 protein is sufficient to activate autophagy. A) Neighbor-joining tree of beclin-1, beclin-1-like, and ATG-6 protein sequences. An alignment of the conserved 17-amino acid portion of beclin-1 is shown at right. B) Predicted 3D- structure (PDB id 4ddp as template) of *Ae. aegypti* ATG-6 (beclin-1-like protein). Orange residues highlight the 17-amino acid peptide used in this study (aa 243-260). C) Schematic representation of full length (FL, upper) and truncated versions *Ae. aegypti* beclin-1 D) Cartoon schematic of ATG-8 expression during autophagy initiation. ATG-8 expression (green) shifts from a diffuse to a punctate pattern as autophagosomes are formed. (E) Merged immunofluorescent staining of ATG-8 (green) in C6/36 cells, with beclin-1-mCherry expression (red) and DAPI-stained cell nuclei (blue). Positive control cells were treated with chloroquine (100 µM) for 6 h; negative control cells were not treated. Beclin-1 treatment groups were transfected with either full-length (FL-beclin-1) or a 17-aa peptide (243-263aa-beclin-1) of *Ae. aegypti* beclin-1 protein. Scale bars=10 µm. Arrowheads mark representative ATG-8 puncta.

### Bioactive peptide of *Ae. aegypti* beclin-1-like induces autophagy *in vitro*

To determine whether the full-length *Ae. aegypti* beclin-1-like protein and/or the conserved 17-aa peptide are sufficient to induce autophagy in mosquito cells, we transfected C6/36 cells to express these proteins (tagged with Flag) using a pAc5-mCherry vector. Specifically, we used plasmids pAc5-AaBecFL-mCh-Flag (full-length beclin-1), pAc5-AaBec243-260-mCh-Flag (17aa peptide of beclin-1), and pAc5-mCherry-Flag (negative control) and screened for markers of autophagy. We used immunofluorescent labeling to screen all treatment groups for ATG8 puncta, which are indicative of autophagy, and found puncta in beclin-1-tranfected cells (full-length and 17-aa peptide) but not in negative controls. Thus we find that both the full-length and 17-aa peptides (AaBec243-260) were sufficient to activate autophagy (Figure 1E).

### *In vitro* characterization of recombinant SINV expressing *Ae. aegypti* beclin-1-like protein

To test the effects of autophagy on SINV replication, we designed a recombinant SINV that expresses AaBec243-260aa peptide during viral replication in cells. Two recombinant SINV strains were generated from a wild-type parental strain (p5’dsMRE16ic, hereafter SINV-WT): p5’dsMRE16ic-AaBec-mCh-Flag (expressing the 17-aa beclin-1 peptide and mCherry, hereafter SINV -AaBec-mCh-Flag), and p5’dsMRE16ic-mCh (negative control expressing mCherry, hereafter SINV-mCh-Flag), (Figure 2A). Each recombinant virus was propagated in BHK-21 cells for seven passages to assess transgene stability on the virus genome. Epifluorescent detection of mCherry expression was confirmed for all passages, and no significant change in the expression of mCherry was noted for any virus (Figure S2). Additionally, the recombinant SINVs were confirmed via RT-PCR using insert-specific primers to produce 648 bp and 573 bp products from recombinant SINV-AaBec-mCh-Flag and recombinant SINV-mCh-Flag, respectively (Figure 2B). No bands of the expected size were observed for the parental wild-type virus.

**Figure 2.**
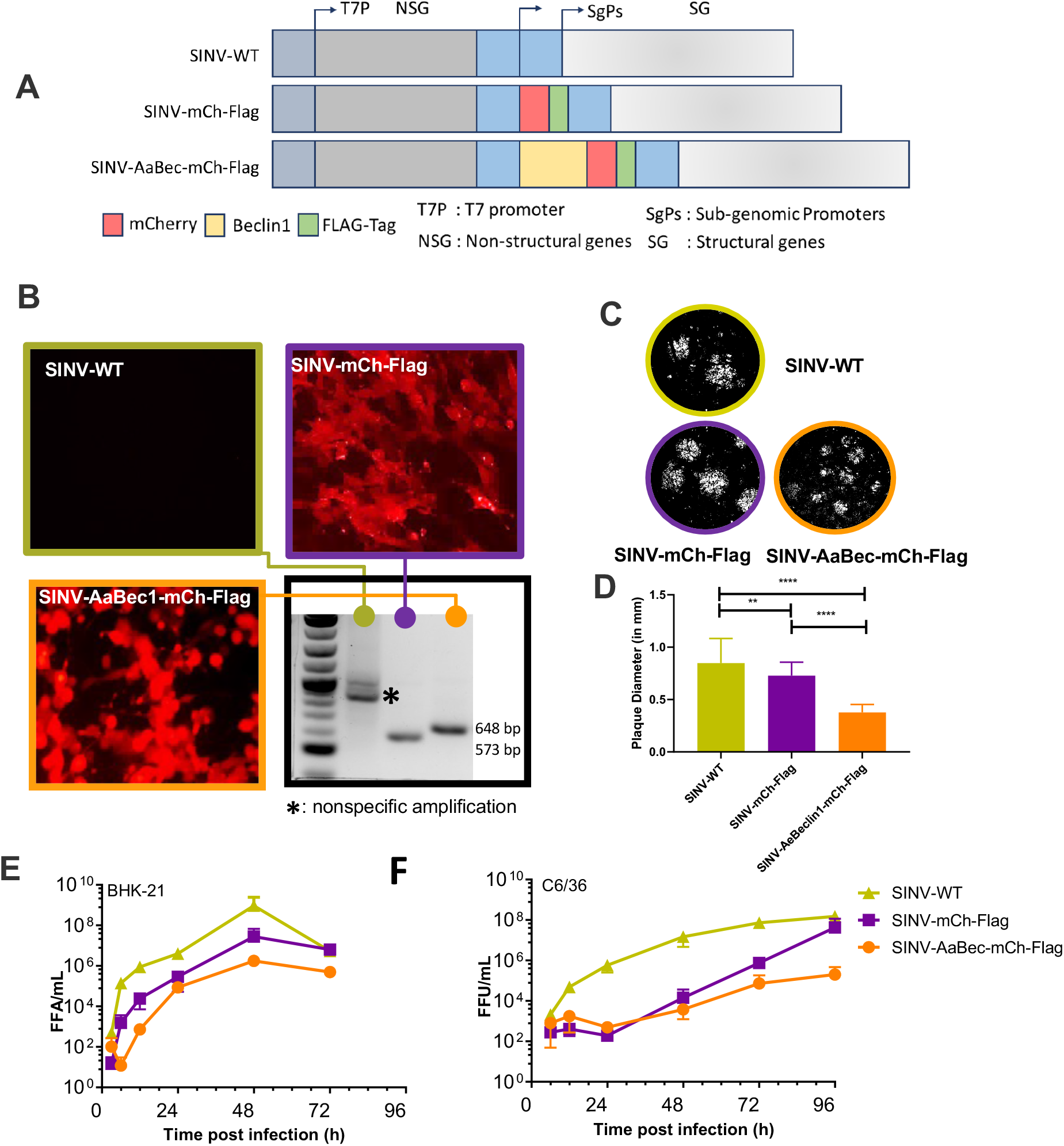
Beclin-1 expressed from recombinant SINV clone affects SINV replication kinetics in vitro. A) Cartoon diagram of recombinant SINV-WT clones, SINV-mCh-Flag and SINV-AaBec- mCh-Flag based on the backbone of 5′dsMRE16ic infectious cDNA clone; constructs express mCherry-Flag and mCherry-beclin-Flag fusion proteins, respectively. The T7 promoter is used to transcribe infectious RNA in vitro. B) Micrographs show mCherry fluorescence in representative cells from each treatment at 24 h post-infection; Agarose gel electrophoresis of RT-PCR amplicons amplified with primers targeted to viral genome flanking the inserts of interest. C) Viral plaque morphology of parental and clone-derived recombinant SINV on Vero cells. Plaques were visualized 72 h post-inoculation by crystal violet staining. D) Plaque diameters for each treatment group. Significance was evaluated by paired t-test. E-F) Replication kinetics of recombinant viruses in BHK21 (left) and C6/36 (right) cells. Infectious virus concentration as FFU/mL in cell supernatants are plotted as a function of time. Each data point represents the log10 mean ± SD FFU/mL from at least two independent tests and error bars show standard deviation. **, P<0.01. ****, P<0.0001. MOI, multiplicity of infection. FFU, focus forming unit. PFU, plaque-forming unit.

We next evaluated plaque size and morphology, indicators of viral growth, in Vero cells 72 h post-infection. Recombinant SINV expressing AaBec-mCh-Flag formed significantly smaller (∼0.3mm) plaques compared to SINV expressing mCherry only (∼7mm) and wild-type SINV (∼8mm) (Figure 2D), indicating slower growth compared to controls.

To understand the role of autophagy in SINV replication, we evaluated viral replication across time for each virus in both C6/36 (mosquito) and BHK-21 (hamster) cells following infection with SINV strains. Cells were infected with the virus at a multiplicity of infection (MOI) of 0.1 and harvested at 6, 12, 24, 48, and 72 h post-infection. An additional time point of 96 h post-infection was analyzed for mosquito cells. SINV expressing SINV-Ae.Bec-mCh-Flag produced >2 logs lower viral titers from 12 h post-infection and longer in both cell lines compared to the SINV-WT. The difference between SINV-Ae.Bec-mCh and SINV-mCh-Flag control was similar in the early time points and showed a difference of ≥ 0.5 logs at 24 h post-infection for BHK-21 and 48 h post-infection for C6/36 cell. Note, in mosquito cells by 96 h post-infection the mCherry control virus titer difference with SINV-Ae.Bec-mCh-Flag reaches a titer similar to the parent wild-type virus. The maximum difference in viral titers between the parent and beclin1-expressing virus was at 48 h post-infection in BHK-21 and 96 h post-infection in C6/36 cells. The average peak titer in BHK-21 was 1.7 × 10^6^ FFU/7.8 × 10^7^ FFU/ml, and 9.3 × 10^8^ for SINV-AaBec-mCherry-Flag, mCherry-Flag only and parent virus, respectively. In cells C6/36 cells, the peak titer of SINV- AaBec-mCh-Flag was 2 × 10^5^ FFU/ml versus 4.2 × 10^7^ FFU/ml in mCherry-Flag control and 1.5 × 10^8^ in the parent virus.

Next, we measured extracellular and intracellular virus particles *in vitro* using the same recombinant viruses. The extracellular infectious virus titer of SINV-AaBec-mCh-Flag at 24 h post-infection showed an ∼ 3.5-log reduction in viral titer in comparison to the parent and SINV- mCh-Flag virus strains (Figure 3A). At 48 h post infection, the same titer difference was reduced to ∼2-log (Figure 3A). A significant difference in viral titer between the mCherry-expressing recombinant virus and parent virus was also noted at both time points (Figure 3A). Intracellular viral titers were even more divergent between groups. Notably, we did not detect any infectious viral particles in the intracellular/cell lysate samples at 24 or 48 h post-infection in AaBec-mCh-Flag expressing cells (Figure 3B). Intracellular titers were high in the control groups, though titers were lower for the mCherry, and Flag tag-expressing recombinant virus compared to parental virus (Figure 3B).

**Figure 3.**
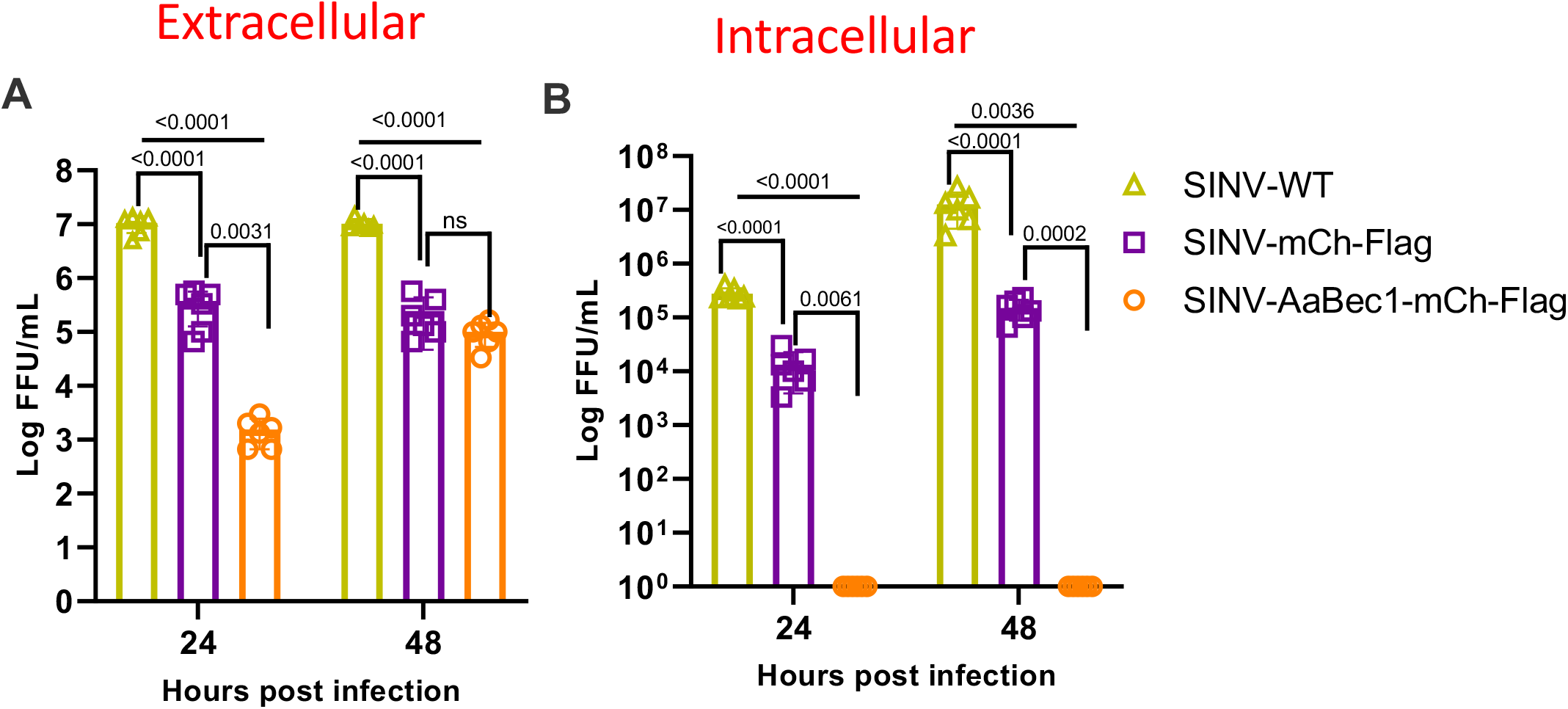
Beclin-1 17aa peptide impairs production of both intracellular and extracellular infectious virus in vitro. C6/36 cells were infected with 0.1 MOI of wild type or recombinant virus as depicted. Extracellular (A) and intracellular (B) progeny viral loads were titrated using focus- forming assays in Vero cells. Means ± SDs from three independent replicate experiments are plotted. t-tests were used to evaluate significance. MOI, multiplicity of infection. FFU, focus forming unit.

### Beclin-1 like protein expression reduces *in vivo* SINV infection rate and viral titer

To examine the effects of 17-aa beclin-1 like bioactive peptide of *Ae. aegypti in vivo*, we intrathoracically injected SINV-WT, SINV-mCh-Flag, and SINV-AaBec-mCh-Flag into adult female *Ae. aegypti*. Following microinjection of 2-3 day old mosquitoes with viruses, we quantified infectious viral titer of mosquitoes at 5 days post infection (dpi) using focus forming assays (FFA). Infections were additionally verified using fluorescence microscopy. By 5 dpi, mosquitoes infected with recombinant viruses (i.e., with mCherry) showed visible red fluorescence, while those infected with wild-type virus did not (Figure 4A-C). We also confirmed the stability of recombinant SINV strains using PCR and fragment analysis (Figure 4D).

**Figure 4.**
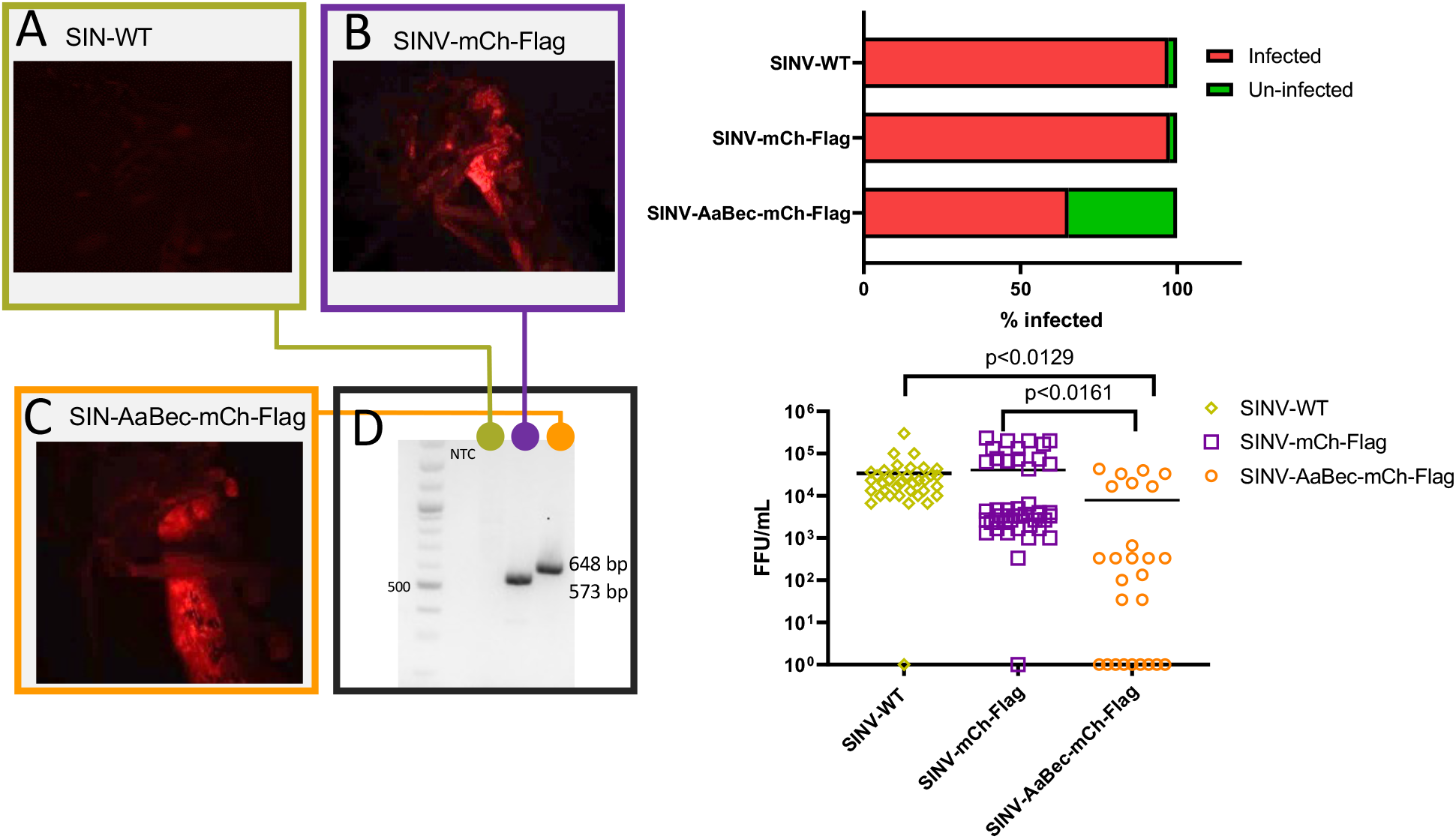
Autophagy negatively impacts SINV infection and viral load in *Ae. aegypti in vivo*. Adult female mosquitoes were intrathoracically micro-injected with SINV-WT, SINV-mCh-Flag, or SINV-AaBec-mCh-Flag. Infections were confirmed by immunofluorescence (A-C) and RT- PCR (D). (E-F) Infection rate and viral loads in whole mosquitoes (expressed as FFU/mL) were analyzed 5 days post injection (dpi). Significance was evaluated using Kruskal Wallis tests.

Interestingly, we found expression of 17-aa beclin-1 peptide suppressed both infection rate as well as SINV viral titer *in vivo*. The infection rates of *Ae* .*aegypti* were significantly lower for beclin-1 expressing SINV (68.38%) in comparison to wild-type (97.22%) and mCherry expressing control (97.72 %) virus (Figure 4E). Further, SINV-AaBec-mCh-Flag titers were significantly lower than those of SINV-WT (*P*<0.0129), and significantly reduced compared with SINV-mCh-Flag (*P*=0.0161). Conversely, there were no significant differences between SINV-WT and SINV- mCh-Flag (Figure 4F).

## Discussion

This study examines the effects of autophagy on arboviral replication in the mosquito *Aedes aegypti*. Using the autophagy inducer ATG6/beclin-1-like protein, we demonstrate negative regulation of SINV both *in vitro* and *in vivo*.

We identified a conserved 17-amino acid peptide of the beclin-1-like protein of *Ae. aegypti* and demonstrated that it is sufficient to activate autophagy in mosquito cells. In C6/36 cells, both full-length beclin-1 and bioactive beclin-1 peptides induce autophagy, confirmed by analyzing cytoplasmic punctate structures. These puncta are a result of the accumulation of lipidated ATG8, ortholog of vertebrate LC3, at phagophores, a characteristic marker of induction of autophagy. ^40^ Similar to a previously characterized bioactive peptide of the human beclin-1 protein, we found the *Ae. aegypti* 17-aa bioactive peptide mapped to the surface of the beclin-1 protein (Figure 1B). ^39^ The human peptide was shown to induce autophagy by binding to the negative regulator of autophagy, Golgi-associated plant pathogenesis-related protein 1 (GAPR-1), and its mosquito ortholog may work via the same mechanism. These studies both demonstrate the potential role of beclin-1 in treating infectious diseases.^39^

Recently, pharmacological modulators such as rapamycin, bafilomycin, and chloroquine have been employed to investigate the role of autophagy in arboviral infection in mosquito cell lines. These studies have produced mixed results. One study reported that autophagy activators restricted SINV replication in C6/36 cells but promoted it in Aag2 cells, while another study showed that autophagy increased Dengue virus titer in Aag2 cells but not in C6/36 cells. ^41, 42^ These reports are challenging to interpret, as the variation in effect may trace back to the different tissue origins of the cell lines tested, or to off-target effects of the pharmacological inhibitors used in those studies, rather than to a mechanism inherent to the autophagy pathway. To avoid pharmacological confounds, here we engineered a recombinant SINV that overexpresses the bioactive beclin-1 peptide of *Ae. aegypti* to selectively activate autophagy. Because this peptide was fused with a fluorescent reporter (mCherry) and a Flag-tag, we also compared it to an mCherry-and Flag-tag only control virus. Our genetic approach using a recombinant virus to express beclin-1 would eliminate any potential off-target effects of exogenous compounds, and is also designed to activate autophagy in the same cells that are infected with the virus. However, one drawback is that the incorporation of the transgene could have destabilized the RNA virus genome. To test this, we propagated the recombinant viruses for seven generations while monitoring fluorescence stability, then verified transgene stability in the genome using RT-PCR (Figure S2). We are, therefore, confident that our experiments are not affected by transgene instability.

Reduced production of recombinant viral progeny when the virus expresses bioactive beclin-1 peptide compared to the mCherry-Flag-only control in C6/36 cells indicates that autophagy negatively regulates SINV. Though untested at this time, autophagy may affect other alphaviruses in the same way. An interesting observation in this experiment was that the mosquito AeBec243- 260 peptide downregulated SINV replication in mammalian cells (BHK-21) although not as strong as the mosquito cell (SINV-mCh-Flag vs SINV-AaBec-mCh-Flag, Fig 2E), suggesting cross-species conservation of function. The strong conservation of peptide residues across this large phylogenetic distance also supports this finding (only 3/17 residues differ between human and *Ae. aegypti*). However, a detailed mechanistic study of this cross-species interaction is beyond the scope of the current study.

The absence of intracellular infectious virus particles in C6/36 cells was surprising (Figure-3). There may be several potential explanations for this observation. One possibility is that autophagy, similar to its role in inhibiting the assembly of Hepatitis B virus capsids, may have played a role in breaking down the intracellular virus maturation vesicles of the SINV, resulting in a significant reduction of SINV assembly and maturation within the cell. ^43^ Another possibility is that, like its effect on the formation and secretion of exosomes, autophagy may have influenced the biogenesis, including the lipid and protein composition of the membrane, transportation, and maturation of intracellular vesicles in cells infected with SINV-AaBec-mCh-Flag. ^44, 45^

There was a broad concordance between our *in vitro* and *in vivo* results. Similar to our *in vitro* findings, the expression of beclin-1 peptide in whole mosquitoes resulted in a decreased infection rate and viral titer, indicating that autophagy negatively regulates the lifecycle of SINV in its invertebrate host. This suggests that autophagy may be a crucial factor in determining vector competence for arbovirus transmission, though further research is needed to confirm this. The mechanism by which autophagy limit viral infections is not fully understood but it may include direct degradation of viral components, or stimulation of adaptive immune responses. Our future studies will aim to identify the interaction partners of beclin-1, understand modulation of invertebrate antiviral responses, and investigate other important components of the autophagy pathway in mosquito-arboviral interactions.

## Material and Methods

### Cells and virus

*Aedes albopictus* C6/36 cells were grown in Schneiders insect cell culture medium (Gibco; 21720024) supplemented with 10% fetal bovine serum (FBS; HyClone, SH 30070). Baby Hamster Kidney (BHK-21) cells and African green monkey kidney (Vero) cells (ATCC) were grown in Eagle’s minimal essential medium (MEM; Gibco) with 10% FBS, 100 µg/ml penicillin/streptomycin and 2 mM L-glutamine and cultured at 37 °C with 5 % CO_2_, and C6/36 cells were maintained at 27 °C. The infectious cDNA clone of SINV, p5’dsMRE16ic-Mx, was kindly gifted by Dr. Clem, Kansas State University. ^46^

### Mosquito colony

*Aedes aegypti* mosquitoes (Rockefeller strain) were obtained from Johns Hopkins University and maintained at 27 °C ± 1 °C, 12:12 h light: dark diurnal cycle at 80% relative humidity in 30×30×30 cm cages. Larvae were fed with ground fish flakes (TetraMin, Melle, Germany), and adult mosquitoes were provided with 10% sucrose solution *ad libitum* for maintenance.

### Expression plasmid construction

Plasmids were generated by Genscript. In brief, pAc5-DmYPss-DmYP1-EGFP was digested with KpnI and HindIII to remove DmYPss-DmYP1. ^47^ *Aedes aegypti* full-length beclin-1-like protein and 243-260 amino acid region fused with mCherry and FLAG tag, from 5’ to 3’ respectively, were synthesized and inserted into the digested backbone to generate pAc5-AaBecFL-mCh-Flag and pAc5-AaBec243-260-mCh-Flag.

### Transfection, chloroquine treatment, and ATG8 immunofluorescent labelling

C6/36 cells were seeded at a density of 1 × 10^6^ cells/well in a 2-well glass chamber slide and allowed to attach overnight. Using lipofectamine 3000, cells were transfected with pAc5- AaBecFL-mCh-Flag and pAc5-AaBec243-260-mCh-Flag constructs. Cells were treated with 100 µM chloroquine for 6 h and untreated cells were used as positive and negative controls, respectively. 48 h post-transfection (or 6 h post-chloroquine treatment), cells were fixed with 4% paraformaldehyde at room temperature and processed for ATG8 staining following standard immunostaining procedure. In brief, fixed cells were permeabilized and blocked for 30 minutes at room temperature with 2% BSA in PBS with 0.01% triton-X. Then the permeabilized cells were incubated overnight at 4 °C with ATG-8 primary antibody reconstituted in permeabilization buffer at 1:200 dilution. Next day, cells were washed and probed with anti-rabbit Alexa-488 secondary antibody and examined under a confocal microscope, and images were processed using ImageJ software.

### Recombinant beclin-1 virus construction

We obtained a gBlock (IDT) of AaBec243-260–mCherry-Flag (bioactive 17 amino acid sequence of the beclin-1-like protein of *Ae. aegypti*) or mCherry only flanked at the 5’ and 3’ end with a 15 nucleotide sequence homologous to the upstream and downstream of the NotI cut site in the p5’dsMRE16ic-Mx construct, respectively. The p5’dsMRE16ic-Mx construct was digested with NotI to remove the Mx gene, and the vector was recovered using standard molecular techniques. The p5’dsMRE16ic-AaBec243-260-mCh-Flag and p5’dsMRE16ic-mCh-Flag was generated using In-Fusion cloning of the recovered vector and the respective gBlocks. The plasmids were verified by Sanger sequencing.

### Virus production

The p5’dsMRE16ic-AaBec243-260-mCh-Flag and the p5’dsMRE16ic-mCh-Flag plasmids were linearized with AscI restriction enzyme and used as templates to generate capped SINV- AaBec243-260-mCh-Flag and SINV-mCh mRNA, respectively, using the SP6 MegaScript (Invitrogen) and m^7^ G(5′)ppp(5′)G Cap Analog (Ambion). BHK-21 cells in 6-well plates were transfected with the capped-SINV-AaBec243-260-mCh-Flag or capped-SINV-mCh-Flag mRNA using Lipofectamine LTX (Invitrogen) and Opti-MEM reduced serum medium (Gibco). Fluorescence was monitored daily for four days. The culture supernatants containing the virus particles were centrifuged at 13,600 x rcf for 5 min to remove cell debris. The supernatant was then aliquoted, stored at -80 °C, and used as seed stock. The recombinant SINV seed stock was further amplified using BHK-21 cells. Recombinant SINV strains were tested for genomic stability by passaging them at least seven times in BHK-21 cells while monitoring fluorescence. The virus stock was titrated using FFAs.

### Plaque forming assays (PFA)

Plaque assays for SINV were performed using Vero cells using standard procedures. ^48^ Briefly, the day before infection, 5×10^5^ Vero cells were seeded on a 6-well plate to achieve a 90-100% confluent cell monolayer. 100 µl of ten-fold serial diluted virus stock/samples were inoculated into Vero cell monolayers and incubated for 1 h at 37 °C and 5% CO_2_ while tilting the plate every 15 min to maintain cell hydration. Unabsorbed virions were washed using DMEM without FBS, then 2 ml of methylcellulose overlay media was added. At 72 h post-infection, the overlay was removed, and cells were fixed and stained with a methanol-crystal violet staining solution. Plaques were counted and imaged. Plaque sizes were measured at the longest diameter using Image J software. Data were analyzed using GraphPad Prism version 8.4.3. paired t- tests were used to compare plaque size across groups.

### Focus forming assays (FFA)

Focus-forming assays for SINV were performed using Vero cells as described previously. ^48^ Briefly, the day before the infection, 3×10^4^ Vero cells were seeded on 96-well plates, resulting in 90-100% confluent cell monolayers. 30 µl of ten-fold serial diluted virus stock/samples were inoculated into Vero cell monolayers and incubated for 1 h in a cell culture incubator at 37 °C and 5% CO_2_. Following adsorption, wells were washed using DMEM without FBS, and 100 µl of methylcellulose overlay media was added. At 24 h post-infection, cells were fixed with 4% paraformaldehyde. Fixed cells were blocked, permeabilized, and probed for SINV viral antigens overnight at 4 °C or 2 h at room temperature using SINV immune ascetic fluid (VR1248AF, ATCC) diluted 1:500 in blocking solution (3% BSA in 1X PBS, 0.1% triton X). Cells were washed three times with PBS and incubated with Alexa Fluor-488 conjugated goat anti-mouse IgG secondary antibody diluted at 1:500 in blocking solution (Invitrogen, USA). Fluorescent foci were visually counted (in a well with a dilution that produced <60 total foci) using an Olympus BX41 microscope with a UPlan FI 4x objective and FITC filter. FFU per ml was calculated by multiplying the number of foci and the dilution factor of the well, then dividing by the inoculation volume (0.03 ml). Data were analyzed using GraphPad Prism version 8.4.3. paired-t tests were used to compare titers across groups.

### Growth kinetics

BHK-21 and C6/36 cell lines were used to assess viral replication. 3×10^4^ cells were seeded in 96- well tissue culture plates and infected with the recombinant virus a day later at an MOI of 0.1. Supernatants were harvested at 24 h intervals, and infectious titers were quantified by FFA on Vero cells.

### Confirmation of recombinant SINV infections in *Ae. aegypti*

*Ae. aegypti* infected with recombinant SINV strains were collected 7 days post-infection, and total RNA was extracted using Direct-zol RNA Miniprep kit (Zymo Research). Reverse transcription PCR (RT-PCR) was performed using the qScript XLT 1-Step RT-PCR kit (Quantabio) and SINV- mChery-Flag-F (CCT-TCA-GCT-TGG-CGG-TCT-G) and SINV-R (TGA-CGC-AGT-AAT-TG- GTG-AGC-G) primer set. The PCR amplicons were electrophoresed and analyzed using a 2% agarose gel.

### Mosquito infections

Adult female *Ae. aegypti* (2–3 days post emergence) were infected with SINV strain MRE-16, and p5’dsMRE16ic-mCh-Flag and p5’dsMRE16ic-AaBec-mCh-Flag by intrathoracic injection. Briefly, each mosquito was cold-anesthetized using a cooling plate and injected with ∼10,000 focus-forming units (FFU) of the selected viral strain in a total volume of 100 nL. Microinjected mosquitoes were placed in cardboard cages with *ad libitum* access to 10% sucrose solution and maintained in the insectary under the same conditions as listed above. After 5 and 7 days post injection (dpi), mosquitoes were sacrificed by overexposing them to triethylamine (Sigma, St. Louis, MO, USA) and placed in 2 mL tubes filled with 1 mL of mosquito diluent (20% heat-inactivated fetal bovine serum (FBS) in Dulbecco’s phosphate-buffered saline (PBS), 50 μg/mL penicillin/streptomycin, 50 μg/mL gentamicin, and 2.5 μg/mL fungizone) and a single zinc-plated steel 4.5-mm bead (Daisy, Rogers, AR, USA). Experimental infections were completed across two replicate batches using the same viral stocks and two subsequent mosquito generations. The results of both replicates were pooled for analysis. A total of 83 mosquitoes were analyzed in the SINV-WT group (46 at 5 dpi and 37 at 7 dpi), 74 mosquitoes in the SINV-mCh-Flag group (46 at 5 dpi and 37 at 7 dpi) and 68 mosquitoes in the SINV-AaBec-mCh-Flag group (26 at 5 dpi and 42 at 7 dpi). Mosquito samples were homogenized at 30 Hz for 2 minutes using a TissueLyser II (QIAGEN GmbH, Hilden, Germany) and centrifuged for 60 s at 11,000 rpm. All samples were further tested for the presence of viral infectious particles by focus forming assay as previously described with one minor modification: viral antigens in infected cells were labeled using Sindbis virus immune ascetic fluid (VR-1248AF; 1:500 dilution). ^49^ Virus titers were calculated and expressed as FFU/mL. Data were analyzed using GraphPad Prism version 8.4.3. Kruskal Wallis tests were used to compare viral titers across mosquito groups.

## Acknowledgements

We would like to thank Drs. Rollie Clem (Kansas State University) and Alexander Raikhel (University of California, Riverside) for providing the p5’dsMRE16ic-Mx Sindbis virus infectious clone and the ATG-8 antibody used in this study. We thank Dr. Hillery Metz for helpful comments on a draft version of this manuscript.

## Funding

This work was funded by NIH grant R01AI128201, USDA Hatch funds (Project #4769), and funds from the Dorothy Foehr Huck and J. Lloyd Huck endowment to JLR, and NIH grant R21AI151475 and a Startup and ASPIRE grant from the University of South Carolina to SP.

## Author contributions

Conceptualization: S.P., J.L.R. Methodology: S.P. C.C.H. Data curation: S.P., C.C.H., M.B., R.M.J. D.K. Funding acquisition: J.L.R., S.P. Writing – first draft: S.P., C.C.H., J.L.R. Writing – review and editing: S.P., C.C.H., M.B., R.M.J, J.L.R. Writing – finalizing MS: S.P, J.L.R.

---

**Figure S1:**
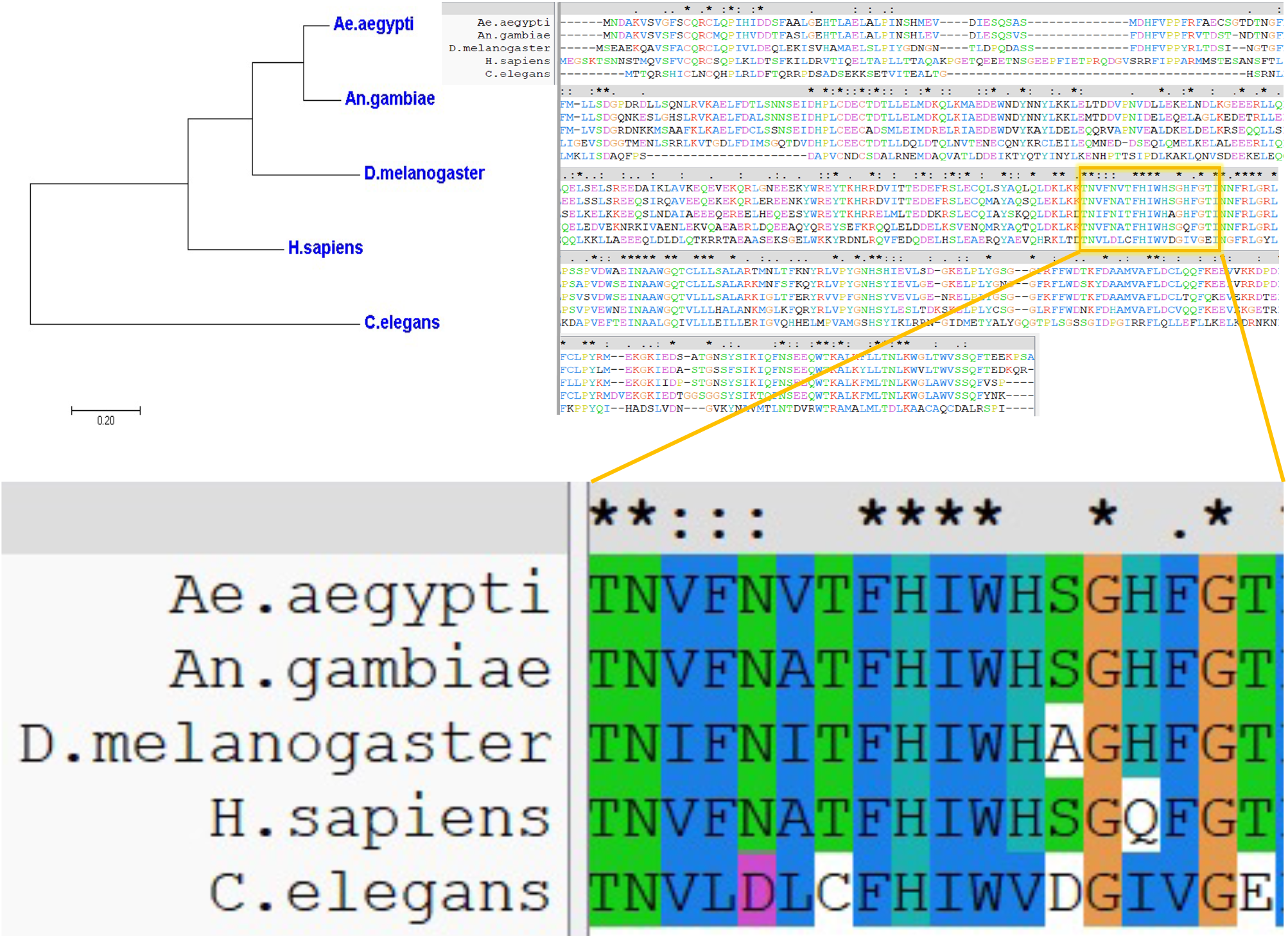
Alignment of *Ae. aegypti* beclin-1 amino acids sequence with orthologs from other species. Enlarged sequence is the minimal peptide used in this study.

**Figure S2:**
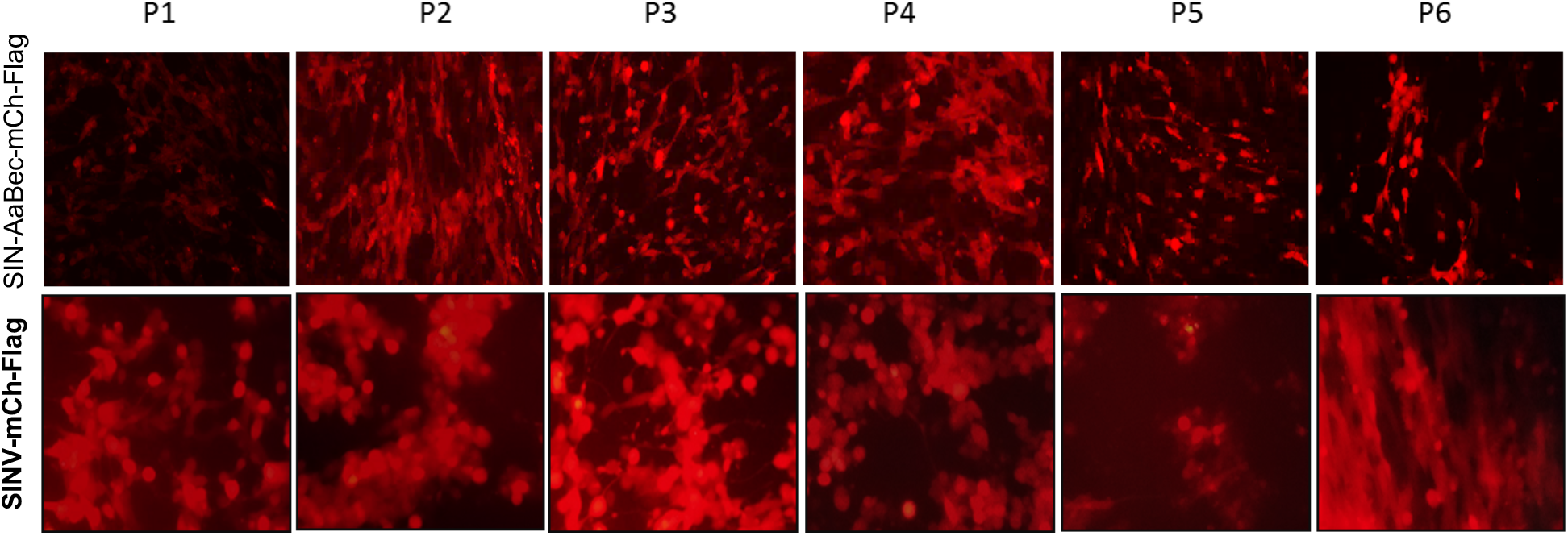
Stability of recombinant SINV. Fluorescence analysis of recombinant SINV expressing mCherry across passages (P1-P6) in cell culture. The recombinant SINV generated by transfecting BHK-21 cells with the transcribed, engineered viral genomic RNA was designated as P0 virus. Virus stocks were passaged seven times (P0−P6) in BHK-21 cells and mCherry expression was visualized with a fluorescence microscope.

## Notes

### Competing Interest Statement

The authors have declared no competing interest.

